# Magnus Representation of Genome Sequences

**DOI:** 10.1101/588582

**Authors:** Chengyuan Wu, Shiquan Ren, Jie Wu, Kelin Xia

## Abstract

We introduce an alignment-free method, the Magnus Representation, to analyze genome sequences. The Magnus Representation captures higher-order information in genome sequences. We combine our approach with the idea of *k*-mers to define an effectively computable Mean Magnus Vector. We perform phylogenetic analysis on three datasets: mosquito-borne viruses, filoviruses, and bacterial genomes. Our results on ebolaviruses are consistent with previous phylogenetic analyses, and confirm the modern viewpoint that the 2014 West African Ebola outbreak likely originated from Central Africa. Our analysis also confirms the close relationship between *Bundibugyo ebolavirus* and *Taï Forest ebolavirus*. For bacterial genomes, our method is able to classify relatively well at the family and genus level, as well as at higher levels such as phylum level. The bacterial genomes are also separated well into Gram-positive and Gram-negative subgroups.

## 1. Introduction

Phylogenetic trees (Fitch and Margoliash, 1967; Robinson and Foulds, 1981) are a type of branching diagram or dendrogram showing the evolutionary relationships among a list of selected biological species, based on similarities in their physical or genetic characteristics. In the field of computational phylogenetics, numerous methods to draw a phylogenetic tree have been devised. In this paper, we are mainly concerned with a class of methods known as distance-matrix methods (Sourdis and Nei, 1988), which rely on a measure of genetic distance between the sequences to draw the phylogenetic tree. Examples of distance-matrix methods include neighbor-joining (Saitou and Nei, 1987) and UPGMA (unweighted pair group method with arithmetic mean) (Gronau and Moran, 2007). A key part of our method involves creating a distance matrix, from which the phylogenetic tree can be drawn using one of the distance-matrix methods.

Alignment methods are often used in phylogenetic analysis to evaluate sequence relatedness (Ortet and Bastien, 2010; Morgenstern et al., 1996; Brudno et al., 2003). Alignment methods usually involve as the first step the alignment of nucleotides in such a way that they are homologous in every position (Castresana, 2000; Abascal et al., 2010; Gatesy et al., 1993). A potential issue in alignment methods is the difficulty of removing divergent regions or gap positions from an alignment. Divergent regions and gaps result in situations where the positional homology cannot be accurately determined (Castresana, 2000). A few methods have been devised to attempt to remove divergent regions or gap positions from an alignment (Castresana, 2000; Fernandes et al., 1993; Rodrigo et al., 1994; Gatesy et al., 1993). For instance, Rodrigo et al. (1994) proposed a method to remove gap positions and adjacent nonidentical positions. Gatesy et al. (1993) proposed a method to remove certain regions of the alignment sensitive to different gap weights.

Alignment-free methods are based on numerical characterizations of genetic sequences, and are a viable alternative to traditional alignment-based methods which are often complicated to compute. Among all alignment-free methods, the k-mer model method may be the most established and best developed (Wen et al., 2014; Chikhi and Medvedev, 2013; Chor et al., 2009). Methods based on k-mer frequency counting are often used to analyze genomes (Kurtz et al., 2008; Liu et al., 2012). One potential disadvantage of the k-mer model is that the relationships between the k-mers of a sequence are often ignored. (Wen et al., 2014). Also, the internal structure within each k-mer (for example its subsequence structure) is often neglected. For instance, AAAAA and CTGAC are both 5-mers but they have very different subsequence structure, i.e., the number of different subsequences of length 1 to 5. In classical k-mer frequency counting, they are both regarded as 1 count of a 5-mer, disregarding the subsequence structure. In our paper, we introduce a method that captures k-mer frequency information as well as the internal subsequence structure information within each k-mer.

In the field of combinatorial group theory, Wilhelm Magnus studied representations of free groups by non-commutative power series (Lyndon and Schupp, 2015). For a free group *F* with basis *x*_1_, …, *x*_*n*_ and a power series ring Π in indeterminates *ξ*_1_, …, *ξ*_*n*_, Magnus showed that the map *μ* : *x*_*i*_ ↦ 1 + *ξ*_*i*_ defines an isomorphism from *F* into the multiplicative group Π^×^ of units in Π. Using concepts from Magnus’ work, we define the Magnus representation and Magnus vector of a DNA/RNA sequence, and apply them in the analysis of genomes (Huang, 2016; Dong et al., 2017; Kwan and Arniker, 2009). In brief, we map a genome sequence *S* = *x*_1_*x*_2_ … *x*_*N*_ of length *N* to the product 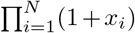, analogous to the map *μ* used by Magnus. For instance, the sequence *AC* will be mapped to (1 + *A*)(1 + *C*). We refer the interested reader who is interested in Magnus’ work on free groups to the book (Lyndon and Schupp, 2015), Chapter I.10, though we emphasize that it is not necessary for a full understanding of our method. In subsequent sections, we will explain our method, with examples, such that it is accessible to readers without a background in abstract algebra.

## 2. Materials and methods

### 2.1. Magnus Representation and Magnus Vector

The Magnus representation and Magnus vector of a DNA/RNA sequence are described as follows. Consider a DNA sequence *S* = *x*_1_*x*_2_ … *x*_*N*_ of length *N*}, where the *x*_*i*_ lie in the set {*A, C, G, T*} (or {*A, C, G, U* in the case of RNA). Our subsequent notation will mainly follow that of DNA sequences for convenience, but we emphasize that our methods work for RNA sequences as well. We define the Magnus representation of *S*, denoted *ρ*(*S*), to be the product 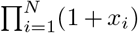 in the non-commutative polynomial algebra *R*⟨*A, C, G, T*⟩, where *R* is a commutative ring. In practice, we may take *R* to be the set of real numbers ℝ, the set of integers ℤ or the ring of integers modulo 2, ℤ/2. In this paper, we use *R* = ℤ (which is equivalent to *R* = ℝ). For most applications, we recommend that users should choose *R* = ℤ. We mention the other commutative rings for completeness, as well as for special cases (for instance the ring *R* = ℤ/2 could be used for coarser classification of genomes).

The Magnus vector of a DNA sequence *S*, denoted by *v*(*S*), is obtained by two steps:

1. Arrange the set of possible words over the alphabet {*A, C, G, T*} of length less than or equal to *N* first by ascending order of length and then by lexicographic order.
2. With respect to the above arrangement, assign *c* ∈ *R* for each term present in *ρ*(*S*) with coefficient *c*, and 0 for each term not present in *ρ*(*S*).

The Magnus vector is a 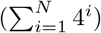-tuple, or equivalently, a 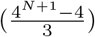-tuple. We illustrate this in the following example, and also summarize it in Table 1. Consider the DNA sequence *S* = *ACT*. Then, the Magnus representation is *ρ*(*S*) = (1 + *A*)(1 + *C*)(1 + *T*) = 1 + *A* + *C* + *T* + *AC* +*AT* + *CT* + *ACT*. The arrangement of the set of possible words of length less than or equal to 3 is: A, C, G, T, AA, AC, AG, AT, CA, CC, CG, CT, GA, GC, GG, GT, TA, TC, TG, TT, AAA, AAC, AAG, AAT, …, TTG, TTT. We note that there are 4^1^ + 4^2^ + 4^3^ = 84 such subsequences. Hence, the Magnus vector is the 84-tuple *v*(*S*) = (1, 1, 0, 1, 0, 1, 0, 1, 0, 0, 0, 1, 0, …, 0, 1, 0, …, 0), where the nonzero entries are in the positions 1, 2, 4, 6, 8, 12 and 28 respectively. Note that due to non-commutativity of the variables, if *S*′ = *CAT*, we observe that *ρ*(*S*′) ≠ *ρ*(*S*) and *v*(*S*′) ≠ *v*(*S*).

**Table 1:**
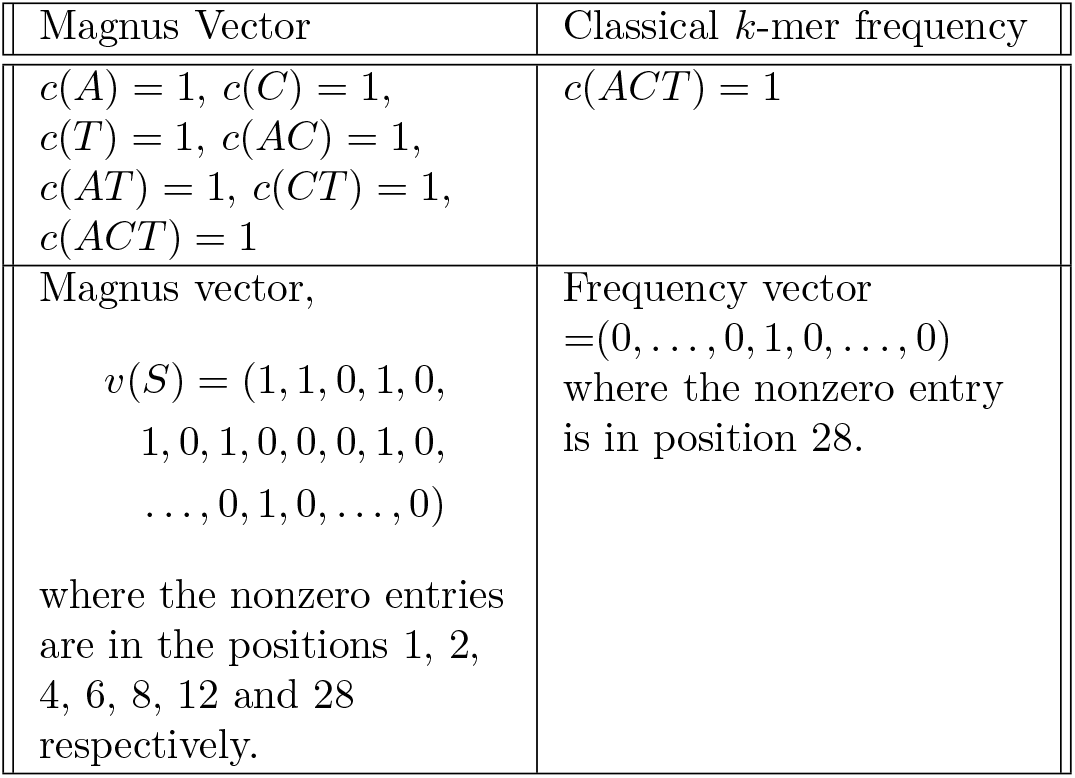
Table illustrating the Magnus vector for the DNA sequence *S* = *ACT*, where *c*(*x*) denotes the function counting the number of occurences of the subsequence *x* in *S*.

| Magnus Vector | Classical $k$-mer frequency |
| --- | --- |
| $c(A) = 1, c(C) = 1,$<br>$c(T) = 1, c(AC) = 1,$<br>$c(AT) = 1, c(CT) = 1,$<br>$c(ACT) = 1$ | $c(ACT) = 1$ |
| Magnus vector,<br>$v(S) = (1, 1, 0, 1, 0,$<br>$1, 0, 1, 0, 0, 0, 1, 0,$<br>$\dots, 0, 1, 0, \dots, 0)$<br>where the nonzero entries<br>are in the positions 1, 2,<br>4, 6, 8, 12 and 28<br>respectively. | Frequency vector<br>$= (0, \dots, 0, 1, 0, \dots, 0)$<br>where the nonzero entry<br>is in position 28. |

We show an example when *R* = ℤ/2. If *S* = *AA*, then *ρ*(*S*) = (1 + *A*)(1 + *A*) = 1 + *AA*. By looking at the word with greatest length in *ρ*(*S*), we observe that it is the DNA sequence itself. Hence, for *R* = ℝ, ℤ, or ℤ/2, the Magnus representation is faithful (namely injective). This means that the Magnus representation is able to distinguish between any two different DNA sequences, and detect all forms of DNA mutations.

It can be seen that the Magnus vector consists of many coordinates even for *N* = 2, however it is typically sparse. Hence, we introduce the short Magnus vector to compress the Magnus vector into fewer coordinates. The short Magnus vector is defined to be the vector whose first coordinate is the dimension of the Magnus vector (i.e. the number of coordinates 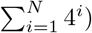, and the subsequent entries indicate the position of the non-zero entries of the Magnus vector. We denote the short Magnus vector also by *v*(*S*), since in practice there is no danger of confusion. In the example of *S* = *AC*, the short Magnus vector is *v*(*S*) = (20, 1, 2, 6). For the example of *S^I^* = *CA*, the short Magnus vector is *v*(*S*′) = (20, 1, 2, 9). If *R* = ℤ/2, there is a one-to-one correspondence between the set of Magnus vectors and the set of short Magnus vectors. In this paper, we do not make use of the short Magnus vector as computationally there is no need to compress the Magnus vector for the value of *k* that we choose (*k* = 5). We mention the short Magnus vector in case users need to use a large *k* for the k-mers, where compression of the Magnus vector may become necessary for computational and data storage reasons.

We remark that there are other theoretical ways to understand the algebra of formal power series on non-commuting variables (see Appendix A).

### 2.2. Summary of the procedures

We compute the mean Magnus vectors of non-overlapping *k*-mers of virus genomes, and their mutual Euclidean distances. We store the distances in a distance matrix and construct a phylogenetic tree (or dendrogram) using neighbor joining or UPGMA.

We choose the neighbor-joining and UPGMA algorithms for several reasons. The primary reason is to correspond to other papers so that we can have a fair comparison. For instance, the two papers Zheng et al. (2015); Li et al. (2017) use the UPGMA algorithm. Since we are using the same virus/bacteria dataset as the two papers, it is more consistent to use the same tree algorithm in Figures 3 and 4 for a fair comparison (with the two papers).

In Figure 2, we use the neighbor-joining algorithm to draw the phylogenetic tree for mosquito-borne viruses. In this case, we are not constrained in our choice of algorithm since we are not following the dataset from any paper. We choose the neighbor-joining algorithm as it is a classical method with a long history of successful applications (Saitou and Nei, 1987; Tamura et al., 2004; Gascuel and Steel, 2006). In addition, by using two different types of trees in our paper, we provide evidence that our method can work on different tree algorithms.

Both neighbor-joining and UPGMA are bottom-up (agglomerative) clustering methods (Gronau and Moran, 2007; Wheeler, 2009). There are some technical differences between the two algorithms: UPGMA assumes a constant rate of evolution for all different lineages, while the neighbor-joining method does not require such an assumption (Kumar et al., 1994). To our knowledge, these technical differences do not make a significant impact on how our method works.

#### 2.2.1. Algorithm for Magnus Vector

We use a quaternary (base 4) system to encode the DNA sequence. Namely, we let the digits 0, 1, 2, 3 represent the letters A, C, G, T respectively. We use the term *DNA subsequence* to denote a sequence that can be derived from the original DNA sequence by deleting some or no letters without changing the order of the remaining letters.

For a DNA sequence of length *N*, we count the number of occurrences of the subsequences “0”, “1”, “2”, “3”, “00”, “01”, etc., up to subsequences of length *N*. The counting can be done efficiently through dynamic programming. For each subsequence *S* (considered as a base 4 number), we convert it to a decimal (base 10) numeral *d*_*S*_. Let *α*_*S*_ be the number of occurrences of the subsequence *S* (in the original DNA sequence), and *l*_*S*_ be the length (number of digits) of *S*. Define 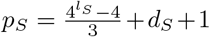. Then, the *p*_*S*_-th component of the Magnus vector is precisely *α*_*S*_.

For instance, consider the DNA sequence CCGAG and the subsequence *S* = *CCG* = 112. Then, *α*_*S*_ = 2, *l*_*S*_ = 3 and *d*_*S*_ = 22. Hence, *p*_*S*_ = 43 and the 43rd component of the Magnus vector is 2.

As an illustration of the algorithm, we list the complete components of the Magnus vector for the example *S* = *AC* = 01. The only subsequences are *S*_1_ = 0, *S*_2_ = 1 and *S*_3_ = 01, each occuring once. Hence, 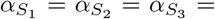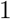. We have 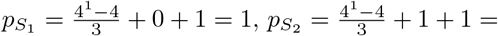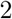, and 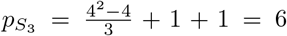. Hence, the complete, components of the Magnus vector in this case is the 20-tuple (1, 1, 0, 0, 0, 1, 0, 0, 0, 0, 0, 0, 0, 0, 0, 0, 0, 0, 0, 0).

We implement the algorithm in Python. The codes in the paper are made publicly available on GitHub (Wu et al., 2019). The codes are also available upon request from the corresponding author.

#### 2.2.2. k-mer and size selection

A *k*-mer is a segment of *k* consecutive nucleotides of a genome sequence (Huang, 2016; Koren et al., 2017; Rizk et al., 2013). Due to the length of the Magnus vector, it is not practical to compute it for the entire genome sequence of length *N*. Instead, we compute the Magnus vector for each *k*-mer, which are enumerated by non-overlapping sliding windows of size *k*, shifting *k* nucleotides each time until the entire sequence (possibly excluding up to *k* − 1 nucleotides at the tail end if *N* is not divisible by *k*) is scanned.

We choose non-overlapping sliding windows due to the observation that the Magnus vectors of overlapping windows may contain similar information (counting the same subsequences). Hence, non-overlapping sliding windows will reduce the total number of windows without much loss of information. The value of *k* is chosen to be 5 as it corresponds to the smallest classification errors of the Baltimore and genus classification labels (Huang, 2016). This leads to the Magnus vector having length 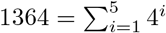.

A schematic diagram is shown in Figure 1. In subsection 6.2, we mention that overlapping *k*-mers is also possible as an improvement (but with increased computational costs).

**Figure 1:**
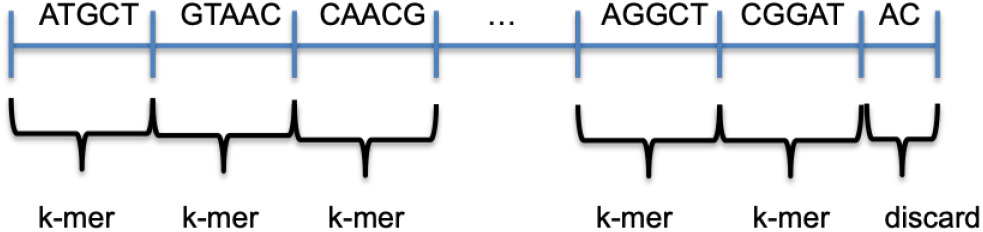
The genome sequence is split into non-overlapping *k*-mers, and the Magnus vector is computed for each *k*-mer, where *k* = 5.

**Figure 2:**
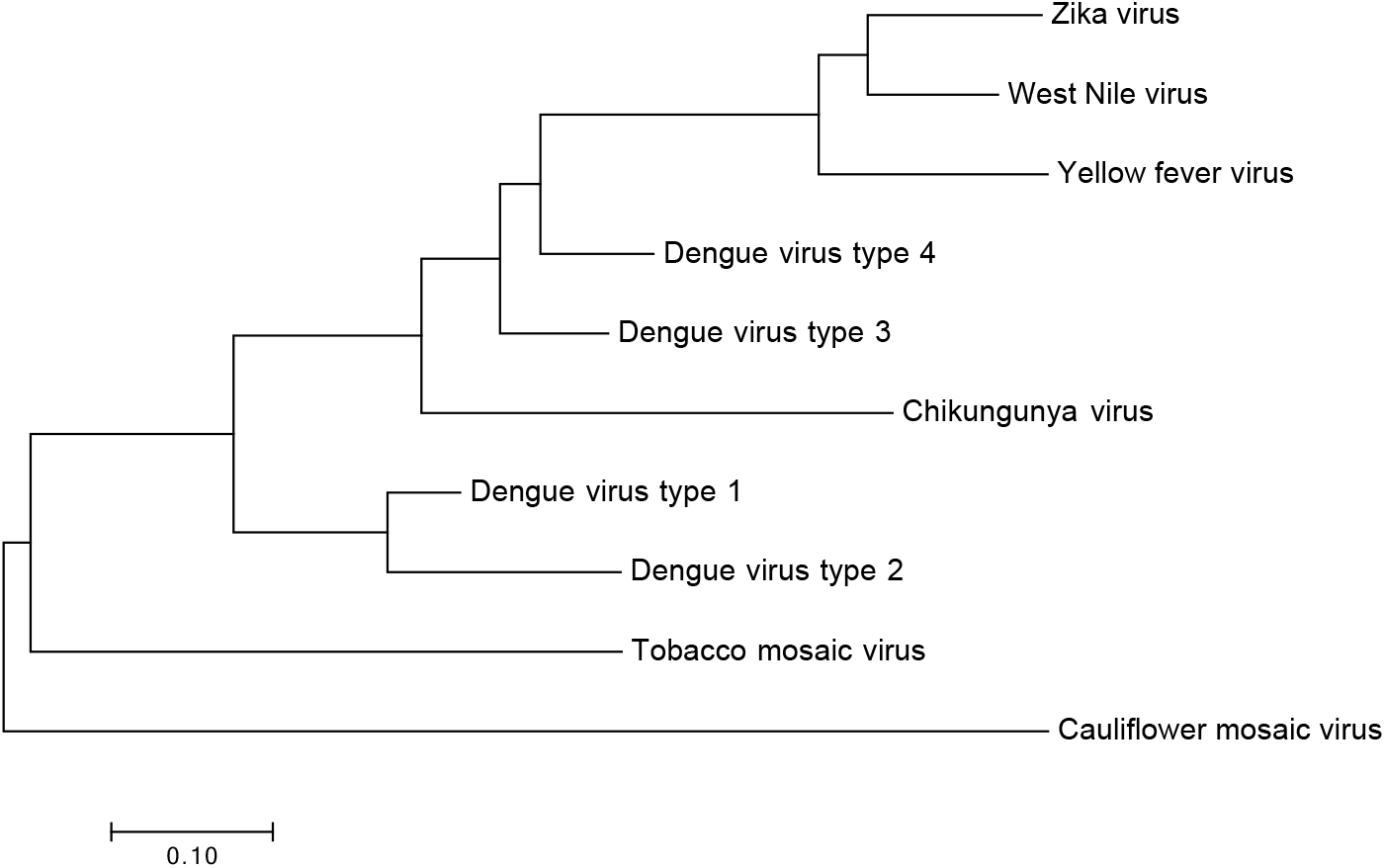
Phylogenetic tree (neighbor-joining) drawn using MEGA7.

**Figure 3:**
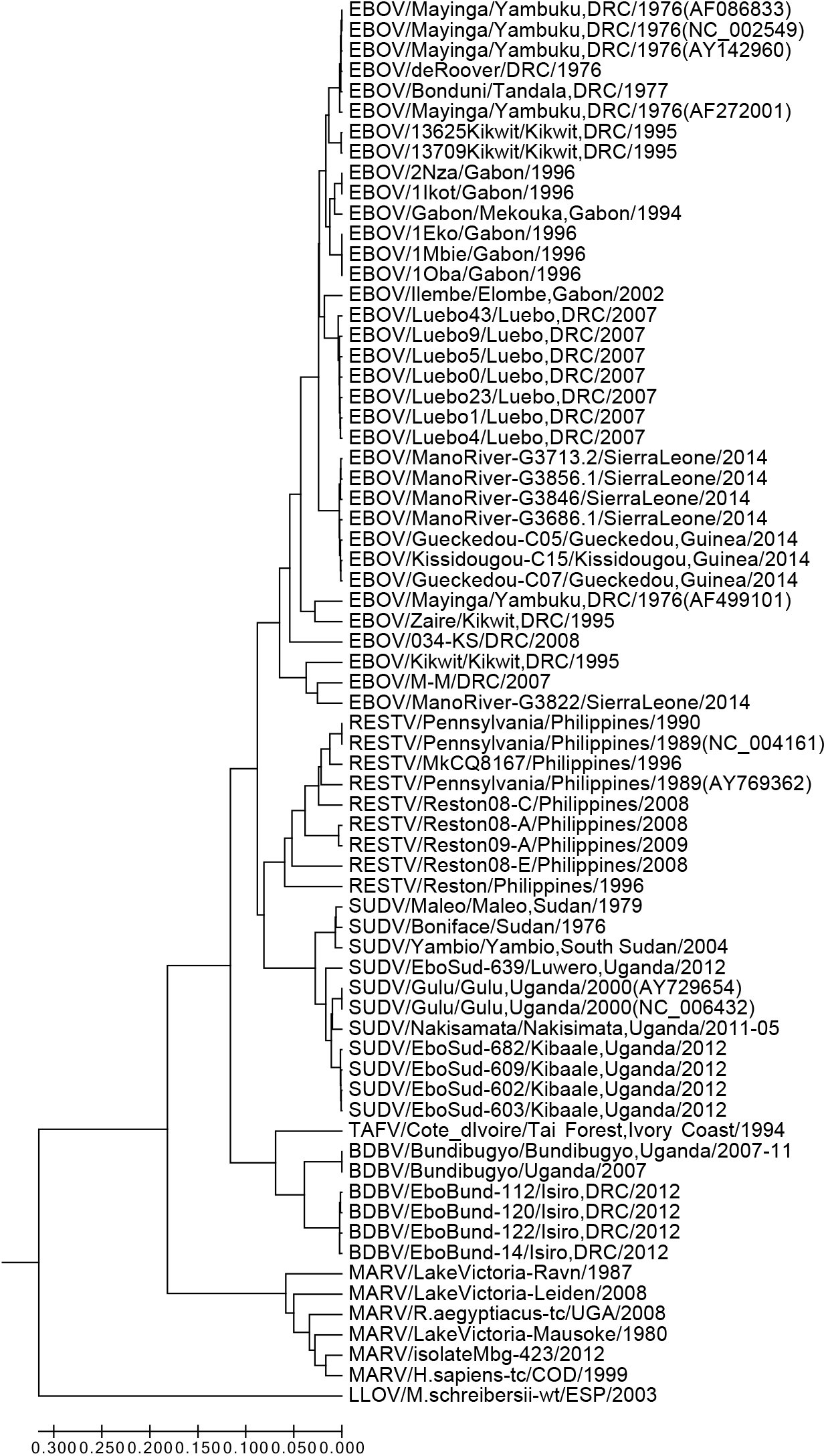
Phylogenetic tree of 69 filoviruses drawn using MEGA7 (UPGMA).

**Figure 4:**
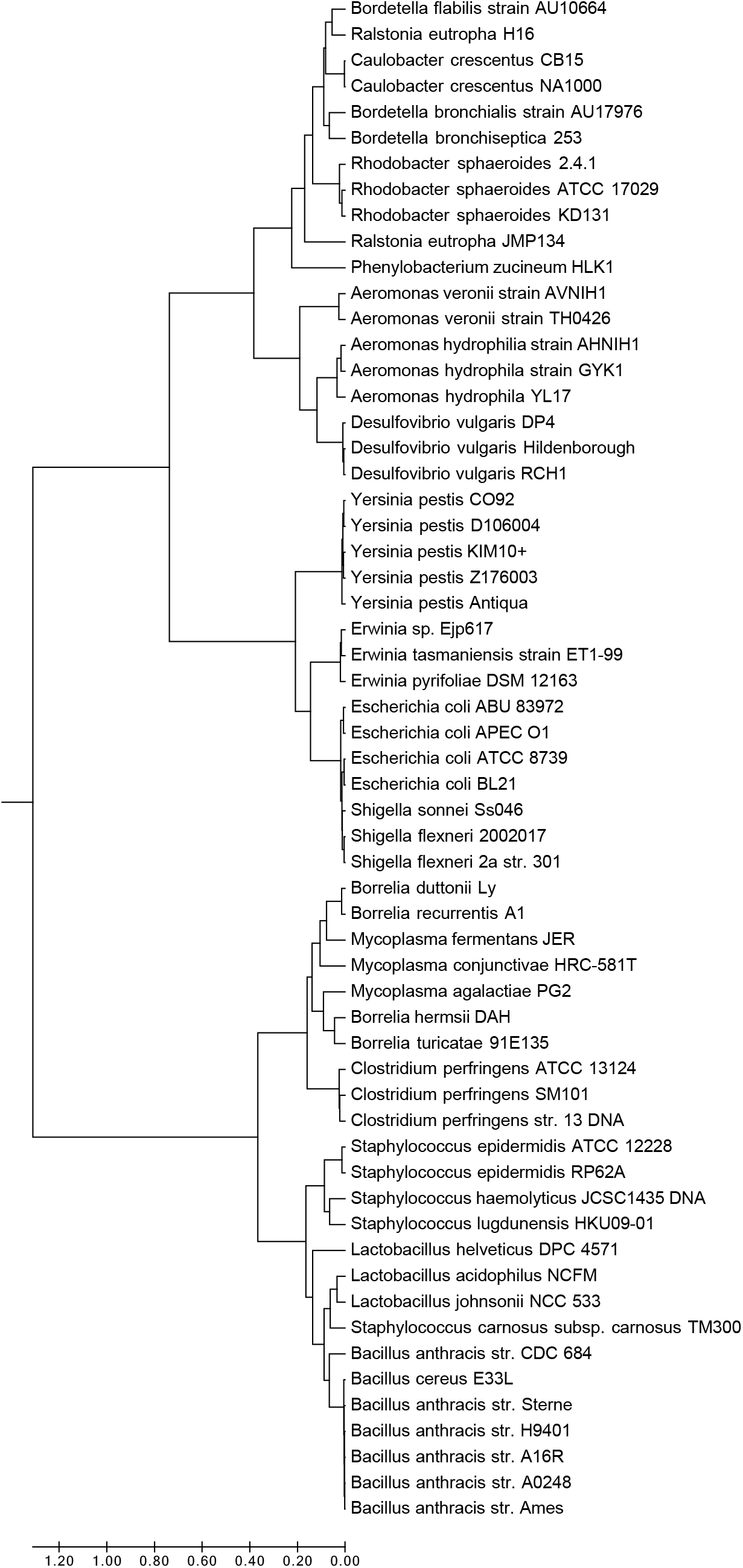
Phylogenetic tree of 59 bacteria drawn using MEGA7 (UPGMA).

#### 2.2.3. Mean Magnus Vector and Euclidean distance

We calculate the *mean Magnus vector* of a genome sequence by dividing the sum of Magnus vectors of all *k*-mers, by the total number of windows. We use the Euclidean distance to calculate the distance between the mean Magnus vector of two different genome sequences.

We use the mean Magnus vector (instead of the Magnus vector) so that we can combine all the information of the Magnus vectors into a single vector representing the genome sequence. For a genome sequence of length *N*, there are 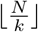 Magnus vectors corresponding to the non-overlapping *k*-mers. Thus, for large *N* we have many Magnus vectors for a single genome sequence. Hence, it is computationally more efficient to combine the Magnus vectors into a single mean Magnus vector which is the representative of the genome sequence. Subsequently, when comparing two genome sequences, we can compare their respective mean Magnus vectors.

#### 2.2.4. Benefits of our approach

The Magnus Representation (and Magnus vector) contains higher-order information about the genome sequence, in the form of its subsequences. This is in contrast to lower-order information such as simply counting the number of letters ‘A’, ‘C’, etc., in the genome sequence. The non-commutativity of the variables enhances the discriminatory power of the Magnus Representation by distinguishing between permutations of subsequences.

By combining the Magnus Representation with the idea of *k*-mers, we improve the computability issue of our approach. (It is infeasible to compute the Magnus Representation of an entire long genome sequence.) By breaking up the genome sequence into *k*-mers, the mean Magnus vector of a large genome sequence can be effectively computed.

### 2.3. Data

#### 2.3.1. Mosquito-borne viruses

We use virus genome data from GenBank (Benson et al., 2008, 2012), with a focus on mosquito-borne viruses such as dengue (Tuiskunen Bäck and Lundkvist, 2013). We also include two plant viruses (*tobacco mosaic virus* and *cauliflower mosaic virus*) for contrast. We compute their mean Magnus vector and the corresponding distance matrix. We then draw a phylogenetic tree (neighbor-joining) using MEGA7 (Kumar et al., 2016).

#### 2.3.2. Ebolaviruses

Following the seminal paper by Hui Zheng, Stephen S.-T. Yau and coauthors (Zheng et al., 2015), we study 69 filoviruses, and draw the phylogenetic tree. Virus genome data are also taken from GenBank. The 69 filoviruses correspond exactly to the ones in Figure 2 and Supplementary Table S2 of Zheng et al. (2015).

#### 2.3.3. Bacterial genomes

To further confirm the advantage of our method, we test bacterial genomes which typically have longer sequences than viruses. Following a landmark study by Li et al. (2017), we analyze a dataset consisting of 59 bacterial species. The genome length in the dataset typically range from 3 to 10 Mb (million base pairs). The 59 bacterial species come from 15 different families, and corresponds to Figure 6 and Table S6 (in the supplementary file) of (Li et al., 2017).

## 3. Results

### 3.1. Mosquito-borne viruses

The phylogenetic tree (strictly speaking, a dendrogram) is drawn using MEGA7 by neighbor-joining and shown in Figure 2.

### 3.2. Ebolaviruses

The phylogenetic tree was drawn using the UPGMA (unweighted pair group method with arithmetic mean) method, using MEGA7. The optimal tree with the sum of branch length = 2.09343667 is shown in Figure 3. The tree is drawn to scale, with branch lengths in the same units as those of the evolutionary distances used to infer the phylogenetic tree. The evolutionary distances were obtained by calculating the Euclidean distance between the respective mean Magnus vectors.

We chose the UPGMA method to correspond to Figure 2 in Zheng et al. (2015).

In the process, we also discover that some viruses are labelled as distinct in GenBank, but are actually the same virus with identical DNA. They are the following pairs of viruses (denoted by their GenBank Accession Number): (FJ217161, NC_014373), (AF522874, NC_004161), (AY729654, NC_006432).

### 3.3. Bacterial genomes

The phylogenetic tree was drawn using the UPGMA method, using MEGA7. The optimal tree with the sum of branch length = 7.27405405 is shown in Figure 4. Similarly, we chose the UPGMA method to correspond to Figure 6 in Li et al. (2017).

### 3.4. Algorithmic complexity and Computation time

For fixed *k*, the time complexity of enumerating and counting all subsequences in each *k*-mer is *O*(1). For a genome sequence of length *N*, the Magnus vector is computed for each of the *N/k k*-mers. Hence, the overall time complexity of the algorithm is *O*(*N*).

For a typical filovirus of around 18000 base pairs (such as 18939 base pairs for the virus KC545393), computation time takes 1 minute and 45 seconds. The algorithm is run on a 2014 model of MacBook Air with 1.4 GHz Intel Core i5 and 4 GB 1600 MHz DDR3.

For bacterial genomes, we run the algorithm on a 2.30GHz Intel Xeon CPU together with a Nvidia Tesla P100 GPU. For a typical bacterial genome of around 5 Mb (5 million base pairs), computation time takes around 3 hours.

## 4. Discussion

### 4.1. Mosquito-borne viruses

Our results obtained by the Magnus vector approach make biological sense. Intuitively, the mosquito-borne viruses should be more or less similar within the group, but should have big differences compared to plant viruses such as *tobacco mosaic virus* and *cauliflower mosaic virus*. Our results clearly reflect this, as shown in the phylogenetic tree in Figure 2.

Our results also show some interesting phenomena among the mosquito-borne viruses. Among the 4 types of dengue viruses, Type 1 and Type 2 appear to be closely related. On the other hand, Type 3 and Type 4 also appear to be closely related, but having some differences from Types 1 and 2. In short, the 4 dengue viruses seem to form two clusters.

*Zika virus* and *West Nile virus* show signs of having close similarities as well. A plausible biological explanation for this may be the fact that both *Zika virus* and *West Nile virus* are believed to originate from the geographical region in or near Uganda (Central East Africa). *Zika virus* was first discovered in the Zika Forest of Uganda in 1947 (Schwartz, 2016). *West Nile virus* was also originally discovered in Uganda in the year 1937 (Johnston and Conly, 2000), in the West Nile area, which gives rise to its name.

### 4.2. Ebolaviruses

Our results are consistent (though not identical) with Zheng et al. (2015), which employs the reliable and highly successful alignment-free natural vector method (Yu et al., 2013).

In particular, our results also have the following properties shared by Zheng et al. (2015): The five species of the Ebolaviruses are separated well (EBOV, SUDV, RESTV, BDBV, TAFV). The viruses from the same country are (generally, with a few exceptions) classified together within each species. The MARV and EBOV genomes are closer than the LLOV and EBOV genomes. In the branch of EBOV, majority (seven out of eight) of the viruses from Guinea and Sierra Leone of the 2014 outbreak are separated from others. *Bundibugyo ebolavirus* and *Taï Forest ebolavirus* are in the same group.

It was considered unusual by Zheng et al. (2015) that *Bundibugyo ebolavirus* and *Taï Forest ebolavirus* are in the same group according to their classification using natural vectors. This is because *Bundibugyo ebolavirus* is a deadly species while the *Taï Forest ebolavirus* is not so deadly. Our results confirm that, though unusual, it seems that the two viruses are indeed closely related. Hence, it appears that the virulence of ebolaviruses may not directly correlate with their genetic proximity.

Our results show that the Reston virus (RESTV) is most closely related to the Sudan virus (SUDV). This result is consistent with previous known phylogenetic analyses (Cantoni et al., 2016).

Our results also show a close similarity between ebolaviruses from the 2014 West Africa Ebola outbreak and ebolaviruses from Central Africa. In particular, the virus KM233096 from Sierra Leone is classified in the same group as HQ613403 from the Democratic Republic of the Congo. This is consistent with the general consensus among experts that the 2014 West African virus likely spread from Central Africa within the past decade (Gire et al., 2014; Alexander et al., 2015).

According to Alexander et al. (2015), the outbreak in Sierra Leone is believed to have started from the introduction of two genetically different viruses from Guinea. Our results are consistent with this, since the virus KM233096 is in a distinct group from the other 4 viruses from Mano River in Sierra Leone.

### 4.3. Bacterial genomes

At the family level, our method separates 10 out of the 15 families of bacteria well: *Aeromonadaceae*, *Bacillaceae*, *Clostridiaceae*, *Desulfovibrionaceae*, *Erwiniaceae*, *Lactobacillaceae*, *Mycoplasmataceae*, *Rhodobacteraceae*, *Yersiniaceae*, *Enterobacteriaceae*. For the remaining 5 families (*Alcaligenaceae*, *Borreliaceae*, *Caulobacteraceae*, *Burkholderiaceae*, *Staphylococcaceae*), the classification is still mostly accurate, however with some exceptions. For instance, 4 out of 5 of the bacteria in the family *Staphylococcaceae* are clustered together, with the exception of *Staphylococcus carnosus subsp. carnosus TM300*. In general, the exceptions are still found very near their actual family.

Our tree (Figure 4) has some advantages at the genus level, as compared to Figure 6 in Li et al. (2017). For example, in our tree the two genera *Escherichia* and *Shigella* are well separated.

Our tree also has some advantages at the phylum level, as compared to Figure 6 in Li et al. (2017). For example, the four families *Bacillaceae*, *Clostridiaceae*, *Lactobacillaceae*, and *Staphylococcaceae* from the phylum *Firmicutes* are clustered together in a major branch of the tree. In Figure 6 of Li et al. (2017), the family *Bacillaceae* is clustered relatively far away from the other three families from *Firmicutes*.

It should be emphasized though, that the method in Li et al. (2017) is very fast (takes minutes to calculate for bacterial genomes) and hence is exceptionally good considering its speed and the fact that Figure 6 of Li et al. (2017) separates all 15 families of bacteria very well.

Our tree also has the property of separating the bacteria into two biologically significant groups: Gram-positive and Gram-negative. The top part of our tree are the Gram-negative bacteria and the bottom part are mostly Gram-positive bacteria (mostly comprised of members of the phylum *Firmicutes*). This property is also shared by the tree drawn by the 9-mer FFP method (Figure S6) of Li et al. (2017), but not shared by Figure 6 of Li et al. (2017) due to the reasons mentioned in the prior paragraph regarding the phylum *Firmicutes*. We note that the family *Mycoplasmataceae* is in the Gram-positive section of our tree (and the tree by the 9-mer FFP method), despite them being Gram-negative. However, there is solid genetic support for the hypothesis that mycoplasmas have evolved from ancestors of Gram-positive bacteria (Razin et al., 1998; Sladek, 1986). Hence, it can be said that the family *Mycoplasmataceae* is positioned in the correct section of our tree.

## 5. Comparison with Common Sequence Analysis Methods

As the method in our paper is new, we outline some comparisons with common sequence analysis methods.

### 5.1. Algorithmic complexity

Firstly, we compare the algorithmic complexity of various methods. We focus on time complexity (rather than space complexity) as that is usually more important in many computational problems in biology (Baichoo and Ouzounis, 2017). We recall that the time complexity of our method for computing the Magnus vector is *O*(*n*), where *n* is the length of the genome sequence (Section 3.4).

The time complexity of the Needleman-Wunsch algorithm (Nordström et al., 2011; Likic, 2008), a type of global pairwise alignment method, is known to be *O*(*mn*) for two sequences of length *m* and *n* (Baichoo and Ouzounis, 2017). The time complexity of the Smith-Waterman algorithm (Farrar, 2006; Pearson, 1991), a type of local alignment method, is also *O*(*mn*) (Baichoo and Ouzounis, 2017).

The time complexity for counting *k*-mer frequencies is *O*(*n*) (Gunewardena, 2014). Our method on Magnus vector is more complicated and costs more time than the *k*-mer frequency counting method (despite having the same time complexity *O*(*n*)). This is primarily due to our method counting more subsequences than just *k*-mers.

### 5.2. Accuracy

Our method is of comparable accuracy to the *k*-mer feature frequency profile (FFP) method, by comparing the tree drawn by the 9-mer FFP method (Figure S6) of Li et al. (2017) with our tree using the same bacterial dataset (Figure 4). Both our tree and the tree drawn by the 9-mer FFP method separate the bacteria well into Gram-positive and Gram-negative subgroups. Also, as discussed extensively in Section 4, our method is of comparable accuracy with the natural vector method (Zheng et al., 2015; Li et al., 2017; Wen et al., 2014), which is a fast, efficient and accurate alignment-free method.

## 6. Conclusion

We outline some further improvements that can be made to our approach. Some of the improvements may lead to increased computational costs.

### 6.1. Multiple-segmented genomes

Our approach can potentially be applied to multiple-segmented genomes, following the examples of the seminal papers by Huang et al. (2014); Yu et al. (2014). In the paper (Huang et al., 2014), the authors introduce a novel approach for comparing multiple-segmented viruses, to excellent effect. In the paper (Yu et al., 2014), the authors apply Lempel–Ziv complexity to define the distance between two nucleic acid sequences. Then, based on this distance, the authors use the Hausdorff distance to make the phylogenetic analysis for multi-segmented viral genomes. This results in a powerful tool for studying the classification of viral genomes. By using the Magnus Vector to define the distance between two nucleic acid sequences, it should be possible to perform an analogous study for multi-segmented viral genomes.

### 6.2. Overlapping windows

Instead of non-overlapping sliding windows, overlapping windows can be used to increase robustness against frameshift errors in genome sequencing.

### 6.3. Increasing window size

Increasing the window size (increasing *k* for the *k*- mers) also has the benefit of increasing robustness against frameshift errors. This is because with a larger window, subsequences will be more likely to stay in the window despite frameshifts, hence minimizing the effect on the mean Magnus vector.

An increase in the window size also allows the Magnus vector to capture more higher-order information in the form of longer subsequences.

### 6.4. Weighted Magnus Vector

The Hadamard product (entrywise product) of a Magnus vector (*m*_1_, …, *m*_*n*_) with a weight vector (*w*_1_, …, *w*_*n*_) produces a weighted Magnus vector (*m*_1_*w*_1_, …, *m*_*n*_*w*_*n*_). This can be used to emphasize certain subsequences by increasing their weight.

## Acknowledgements

The project was supported in part by the Singapore Ministry of Education research grant (AcRF Tier 1 WBS No. R-146-000-222-112). The first author was supported in part by the President’s Graduate Fellowship of National University of Singapore. The fourth author was supported by Nanyang Technological University Startup Grants M4081842, Singapore Ministry of Education Academic Research Fund Tier 1 RG31/18, Tier 2 MOE2018-T2-1-033.

We wish to thank the referees most warmly for numerous suggestions that have improved the exposition of this paper.

**Appendix A. Supplementary data and proofs**

**References**

## APPENDIX A

## 1. Algebra of formal power series on non-commuting variables

Let *F* be a free group on letters *x*_1_, *x*_2_, …. Let *R* be a commutative ring. Consider the group ring *R*(*F*), which is an algebra over *R*. Consider the augmentation ideal filtration of *R*(*F*), where *IF* = ker(*ϵ* : *R*(*F*) → *R*) with *ϵ*(*g*) = 1 for *g* ∈ *F*. The augmentation ideal filtration is given by *I*^*n*^*F* = (*IF*) · (*IF*) … (*IF*), the *n*-fold product of *IF*. Let 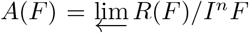, the inverse limit. Then, one can prove that *A*(*F*) is the algebra of formal power series on non-commuting variables *x*_1_, *x*_2_, … using the property that *F* is a free group. The mapping *F* → *A*(*F*) is exactly the Magnus representation. We will describe it in greater detail in the following paragraphs.

### Definition 1.1

(cf. [1, p. 132]). The homomorphism *ϵ* : *R*(*F*) → *R* given by

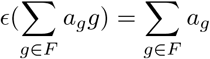

is called the *augmentation mapping* of *R*(*F*) and its kernel, denoted by *IF*, is called the *augmentation ideal* of *R*(*F*).

### Proposition 1.2

(cf. [1, p. 133]). The set {*g* − 1| *g* ∈ *F*, *g* ≠ 1} is a basis of *IF* over *R*.

Thus, we can write

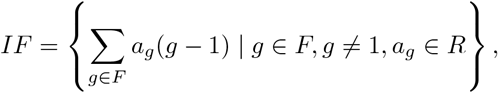

where the sums are finite sums.

*Proof.* If α = ∑_*g*∈*F*_ *a*_*g*_*g* belongs to *IF*, then *ϵ*(α) = ∑_*g*∈*F*_ *a*_*g*_ = 0. Hence, *α* can be expressed in the form:

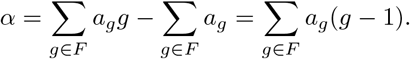

We note that *ϵ*(*g* − 1) = 0 so that *g* − 1 ∈ *IF*. Hence, this implies that {*g* − 1 | *g* ∈ *F, g* ≠ 1} is a generating set for *IF* over *R*. If *a*_1_(*g*_1_ − 1) + · · · + *a*_*n*_(*g*_*n*_ − 1) = 0 for distinct *g*_*i*_ ≠ 1, then *a*_1_*g*_1_ + · · · + *a*_*n*_*g*_*n*_ − (*a*_1_ + · · · + *a*_*n*_)1 = 0. Since *g*_*i*_ ≠ 1, hence we have *a*_1_ = · · · = *a*_*n*_ = 0. This shows linear independence of the generating set. Thus, we have shown that {*g* − 1 |*g* ∈ *F, g* ≠ 1} is a basis for *IF*.

### Corollary 1.3.

The augmentation ideal *IF* is generated by

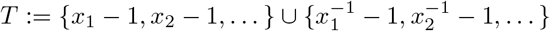

as an *R*-algebra.

That is, every element *α* ∈ *IF* can be expressed as a polynomial with indeter-minates in *T* and coefficients in *R*.

*Proof.* By Proposition 1.2, any element in *IF* is of the form *α* = ∑_*g*∈*F*_ *a*_*g*_(*g* − 1). Since *F* is a free group, any element *g* ∈ *F* is a word in *T*. By applying repeatedly the identity

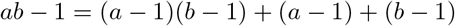

for *a, b* ∈ *F*, we see that *g* − 1 is a polynomial with indeterminates in *T* and coefficients in *R*. Hence, it follows that the same is true for *α*.

Since 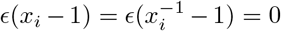, it is clear that conversely, any polynomial with indeterminates in *T* and coefficients in *R* is in *IF*

### Corollary 1.4.

The augmentation ideal *IF* is generated by

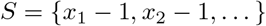

as an ideal of *R*(*F*), or equivalently, as an *R*(*F*)-submodule.

*Proof.* By Proposition 1.2, any element *α* ∈ *F* is of the form *α* = ∑_*g*∈*F*_ *a*_*g*_(*g* − 1), where *g* ∈ *F* is a word in *S*.

By the identities

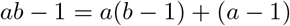

and

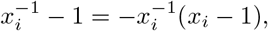

it follows that *g* − 1 is a finite linear combination of {*x*_1_ − 1, *x*_2_ − 1, …} with coefficients in *R*(*F*). Hence, the same is true for *α*.

Conversely, since *ϵ*(*x*_*i*_ − 1) = 0 and *ϵ* is an *R*-algebra homomorphism, it is clear that any linear combination of {*x*_1_ − 1, *x*_2_ − 1, …} with coefficients in *R*(*F*) is in *IF*.

### Definition 1.5

(cf. [2, p. 651]). An *inverse system* in the category of *R*-algebras **R**-**Alg**consists of an ordered pair 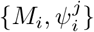, where (*M*_*i*_)_*i*∈*I*_ is a family of *R*-algebras indexed by a partially ordered set (*I*, ⪯) and 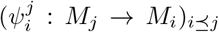 is a family of *R*-algebra homomorphisms, such that the following diagram commutes whenever *i* ⪯ *j* ⪯ *k*:

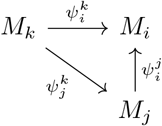

### Proposition 1.6

(cf. [2, p. 652]). For *m* ≥ *n*, define 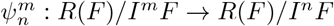 by

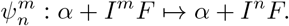

Then, 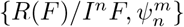 is an inverse system over ℕ.

*Proof. IF* is the kernel of ∈ and thus a subalgebra of *R*(*F*). Each *I*^*n*^*F* is also a subalgebra and there is a decreasing filtration

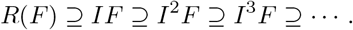

Since *I*^*m*^*F* ⊆ *I*^*n*^*F* for *m* ≥ *n*, the maps 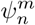 are well-defined. For α + *I*^*k*^*F* ∈ *R*(*F*)/*I*^*k*^*F R*(*F*)/*IkF*, we have

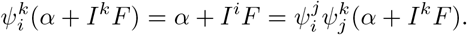

Hence, 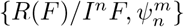 is an inverse system over ℕ.

The categorical definition for inverse limit is stated in [2, p. 653]. The inverse limit is unique up to the isomorphism, if it exists.

### Proposition 1.7

(cf. [2, p. 669]). The inverse limit of an inverse system 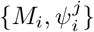 of *R*-algebras over a partially ordered index set *I* exists.

In particular,

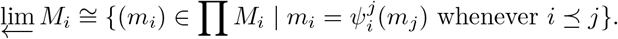

*Proof.* The proof is similar to that of [2, p. 669].

### Theorem 1.8.

Let

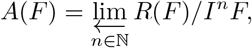

where *F* is a free group with free generating set {*x*_1_, *x*_2_, …}.

Then,

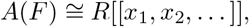

the algebra of formal power series on non-commuting variables *x*_1_, *x*_2_, · · ·.

*Proof.* First, we observe that a change of variables *y*_*i*_ − 1 = *x*_*i*_ induces an isomorphism

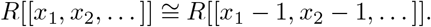

We can further view *R*[[*x*_1_ − 1, *x*_2_ − 1, …]] as the inverse limit

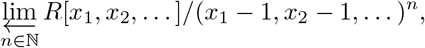

where (*x*_1_ − 1, *x*_2_ − 1, …) denotes the ideal of *R*[*x*_1_, *x*_2_, …] generated by {*x*_1_ − 1, *x*_2_ − 1, …}.

Alternatively, we can view *R*[[*x*_1_ − 1, *x*_2_ − 1, …]] as the inverse limit

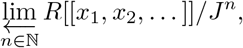

where *J* denotes the ideal of *R*[[*x*_1_, *x*_2_, …]] generated by {*x*_1_ − 1, *x*_2_ − 1, …}.

Consider the homomorphism *ϕ* : *R*(*F*) → *R*[[*x*_*1*_ − 1, *x*_2_ − 1, …]] defined on the generators of *F* by *x*_*i*_ ↦ *x*_*i*_ and extending to *R*(*F*). The map is well-defined because

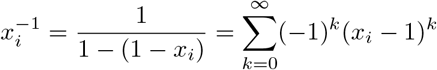

lies in *R*[[*x*_1_ − 1, *x*_2_ − 1, …]].

Let *α* ∈ *IF*. By Corollary 1.3, α can be expressed as a polynomial with indeter-minates in

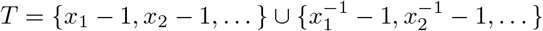

and coefficients in *R*. By the identity

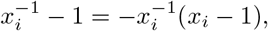

we see that *ϕ*(*α*) ∈ *J*. Hence, we have *ϕ*(*IF*) ⊆ *J*. Similarly, we have *ϕ*(*I*^*n*^*F*) ⊆ *J*^*n*^, for all *n* ∈ ℕ.

Hence, the homomorphism *θ* : *R*(*F*)*/I*^*n*^*F* → *R*[[*x*_1_, *x*_2_, …]]/*J*^*n*^ defined by *θ*(α + *I*^*n*^*F*) = α + *J*^*n*^ is well-defined. The homomorphism then extends to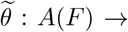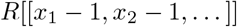

By Corollary 1.4, we see that (*x*_1_ − 1, *x*_2_, …)^*n*^ ⊆ *I*^*n*^*F*. Hence, the map *ψ*: *R*[*x*_1_, *x*_2_, …]/(*x*_1_ − 1, *x*_2_ − 1, …)^*n*^ → *R*(^*F*^)/*I*^*n*^*F* defined by

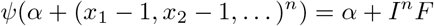

is well-defined. Subsequently,*ψ* extends to 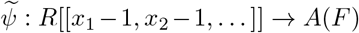 which is the inverse of 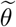. Hence, 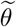 is an isomorphism, and this completes the proof.

We then have the following corollary.

### Corollary 1.9

The mapping *F* → *A*(*F*) defined on the generators by *x*_*i*_ ↦ 1 + *x*_*i*_ and extended to *F* is exactly the Magnus representation.

## References

Abascal, F., Zardoya, R., Telford, M.J., 2010. TranslatorX: multiple alignment of nucleotide sequences guided by amino acid translations. Nucleic acids research 38, W7–W13.

Alexander, K.A., Sanderson, C.E., Marathe, M., Lewis, B.L., Rivers, C.M., Shaman, J., Drake, J.M., Lofgren, E., Dato, V.M., Eisenberg, M.C., et al., 2015. What factors might have led to the emergence of Ebola in West Africa? PLoS neglected tropical diseases 9, e0003652.

Baichoo, S., Ouzounis, C.A., 2017. Computational complexity of algorithms for sequence comparison, short-read assembly and genome alignment. Biosystems 156, 72–85.

Benson, D.A., Cavanaugh, M., Clark, K., Karsch-Mizrachi, I., Lipman, D.J., Ostell, J., Sayers, E.W., 2012. Genbank. Nucleic acids research 41, D36–D42.

Benson, D.A., Karsch-Mizrachi, I., Lipman, D.J., Ostell, J., Wheeler, D.L., 2008. Genbank. Nucleic acids research 36, D25.

Brudno, M., Do, C.B., Cooper, G.M., Kim, M.F., Davydov, E., Green, E.D., Sidow, A., Batzoglou, S., Program, N.C.S., et al., 2003. LAGAN and multi-LAGAN: efficient tools for large-scale multiple alignment of genomic DNA. Genome research 13, 721–731.

Cantoni, D., Hamlet, A., Michaelis, M., Wass, M.N., Rossman, J.S., 2016. Risks posed by Reston, the forgotten ebolavirus. mSphere 1, e00322–16.

Castresana, J., 2000. Selection of conserved blocks from multiple alignments for their use in phylogenetic analysis. Molecular biology and evolution 17, 540–552.

Chikhi, R., Medvedev, P., 2013. Informed and automated k-mer size selection for genome assembly. Bioinformatics 30, 31–37.

Chor, B., Horn, D., Goldman, N., Levy, Y., Massingham, T., 2009. Genomic DNA k-mer spectra: models and modalities. Genome biology 10, R108.

Dong, R., Zheng, H., Tian, K., Yau, S.C., Mao, W., Yu, W., Yin, C., Yu, C., He, R.L., Yang, J., Yau, S.S., 2017. Virus database and online inquiry system based on natural vectors. Evolutionary Bioinformatics 13, 1176934317746667.

Farrar, M., 2006. Striped Smith–Waterman speeds database searches six times over other SIMD implementations. Bioinformatics 23, 156–161.

Fernandes, A.P., Nelson, K., Beverley, S.M., 1993. Evolution of nuclear ribosomal RNAs in kinetoplastid protozoa: perspectives on the age and origins of parasitism. Proceedings of the National Academy of Sciences 90, 11608–11612.

Fitch, W.M., Margoliash, E., 1967. Construction of phylogenetic trees. Science 155, 279–284.

Gascuel, O., Steel, M., 2006. Neighbor-joining revealed. Molecular biology and evolution 23, 1997–2000.

Gatesy, J., DeSalle, R., Wheeler, W., 1993. Alignment-ambiguous nucleotide sites and the exclusion of systematic data. Molecular phylogenetics and evolution 2, 152–157.

Gire, S.K., Goba, A., Andersen, K.G., Sealfon, R.S., Park, D.J., Kanneh, L., Jalloh, S., Momoh, M., Fullah, M., Dudas, G., et al., 2014. Genomic surveillance elucidates ebola virus origin and transmission during the 2014 outbreak. Science 345, 1369–1372.

Gronau, I., Moran, S., 2007. Optimal implementations of upgma and other common clustering algorithms. Information Processing Letters 104, 205–210.

Gunewardena, S.S., 2014. Optimum-time, optimum-space, algorithms for k-mer analysis of whole genome sequences. Journal of Bioinformatics and Comparative Genomics 1, 1.

Huang, H.H., 2016. An ensemble distance measure of k-mer and natural vector for the phylogenetic analysis of multiple-segmented viruses. Journal of theoretical biology 398, 136–144.

Huang, H.H., Yu, C., Zheng, H., Hernandez, T., Yau, S.C., He, R.L., Yang, J., Yau, S.S.T., 2014. Global comparison of multiple-segmented viruses in 12-dimensional genome space. Molecular phylogenetics and evolution 81, 29–36.

Johnston, B.L., Conly, J.M., 2000. West Nile virus-where did it come from and where might it go? Canadian Journal of Infectious Diseases and Medical Microbiology 11, 175–178.

Koren, S., Walenz, B.P., Berlin, K., Miller, J.R., Bergman, N.H., Phillippy, A.M., 2017. Canu: scalable and accurate long-read assembly via adaptive k-mer weighting and repeat separation. Genome research, gr–215087.

Kumar, S., Stecher, G., Tamura, K., 2016. MEGA7: molecular evolutionary genetics analysis version 7.0 for bigger datasets. Molecular biology and evolution 33, 1870–1874.

Kumar, S., Tamura, K., Nei, M., 1994. MEGA: molecular evolutionary genetics analysis software for microcomputers. Bioinformatics 10, 189–191.

Kurtz, S., Narechania, A., Stein, J.C., Ware, D., 2008. A new method to compute K-mer frequencies and its application to annotate large repetitive plant genomes. BMC genomics 9, 517.

Kwan, H.K., Arniker, S.B., 2009. Numerical representation of dna sequences, in: Electro/Information Technology, 2009. eit’09. IEEE International Conference on, IEEE. pp. 307–310.

Li, Y., He, L., He, R.L., Yau, S.S.T., 2017. A novel fast vector method for genetic sequence comparison. Scientific reports 7, 12226.

Likic, V., 2008. The Needleman-Wunsch algorithm for sequence alignment. Lecture given at the 7th Melbourne Bioinformatics Course, Bi021 Molecular Science and Biotechnology Institute, University of Melbourne, 1–46.

Liu, B., Yuan, J., Yiu, S.M., Li, Z., Xie, Y., Chen, Y., Shi, Y., Zhang, H., Li, Y., Lam, T.W., et al., 2012. Cope: an accurate k-mer-based pair-end reads connection tool to facilitate genome assembly. Bioinformatics 28, 2870–2874.

Lyndon, R.C., Schupp, P.E., 2015. Combinatorial group theory. Springer.

Morgenstern, B., Dress, A., Werner, T., 1996. Multiple DNA and protein sequence alignment based on segment-to-segment comparison. Proceedings of the National Academy of Sciences 93, 12098–12103.

Nordström, K.J., Sällman Almén, M., Edstam, M.M., Fredriksson, R., Schiöth, H.B., 2011. Independent HHsearch, Needleman–Wunsch-based, and motif analyses reveal the overall hierarchy for most of the G protein-coupled receptor families. Molecular biology and evolution 28, 2471–2480.

Ortet, P., Bastien, O., 2010. Where does the alignment score distribution shape come from? Evolutionary Bioinformatics 6, EBO–S5875.

Pearson, W.R., 1991. Searching protein sequence libraries: comparison of the sensitivity and selectivity of the Smith-Waterman and FASTA algorithms. Genomics 11, 635–650.

Razin, S., Yogev, D., Naot, Y., 1998. Molecular biology and pathogenicity of mycoplasmas. Microbiol. Mol. Biol. Rev. 62, 1094–1156.

Rizk, G., Lavenier, D., Chikhi, R., 2013. Dsk: k-mer counting with very low memory usage. Bioinformatics 29, 652–653.

Robinson, D.F., Foulds, L.R., 1981. Comparison of phylogenetic trees. Mathematical biosciences 53, 131–147.

Rodrigo, A.G., Bergquist, P.R., Bergquist, P.L., 1994. Inadequate support for an evolutionary link between the Metazoa and the Fungi. Systematic Biology 43, 578–584.

Saitou, N., Nei, M., 1987. The neighbor-joining method: a new method for reconstructing phylogenetic trees. Molecular biology and evolution 4, 406–425.

Schwartz, D.A., 2016. The origins and emergence of Zika virus, the newest TORCH infection: what’s old is new again. Archives of pathology & laboratory medicine 141, 18–25.

Sladek, T.L., 1986. A hypothesis for the mechanism of mycoplasma evolution. Journal of theoretical biology 120, 457–465.

Sourdis, J., Nei, M., 1988. Relative efficiencies of the maximum parsimony and distance-matrix methods in obtaining the correct phylogenetic tree. Molecular biology and evolution 5, 298–311.

Tamura, K., Nei, M., Kumar, S., 2004. Prospects for inferring very large phylogenies by using the neighbor-joining method. Proceedings of the National Academy of Sciences 101, 11030–11035.

Tuiskunen Bäck, A., Lundkvist, Å., 2013. Dengue viruses–an overview. Infection ecology & epidemiology 3, 19839.

Wen, J., Chan, R.H., Yau, S.C., He, R.L., Yau, S.S., 2014. K-mer natural vector and its application to the phylogenetic analysis of genetic sequences. Gene 546, 25–34.

Wheeler, T.J., 2009. Large-scale neighbor-joining with ninja, in: International Workshop on Algorithms in Bioinformatics, Springer. pp. 375–389.

Wu, C., Ren, S., Wu, J., Xia, K., 2019. Magnus-representation. https://github.com/wuchengyuan88/Magnus-Representation.

Yu, C., He, R.L., Yau, S.S.T., 2014. Viral genome phylogeny based on Lempel–Ziv complexity and Hausdorff distance. Journal of theoretical biology 348, 12–20.

Yu, C., Hernandez, T., Zheng, H., Yau, S.C., Huang, H.H., He, R.L., Yang, J., Yau, S.S.T., 2013. Real time classification of viruses in 12 dimensions. PloS one 8, e64328.

Zheng, H., Yin, C., Hoang, T., He, R.L., Yang, J., Yau, S.S.T., 2015. Ebolavirus classification based on natural vectors. DNA and cell biology 34, 418–428.

## References

1. César Polcino Milies and Sudarshan K. Sehgal, An introduction to group rings, vol. 1, Springer Science & Business Media, 2002.

2. Joseph J. Rotman, Advanced modern algebra: Part 1, vol. 165, American Mathematical Soc., 2015.

